# Evidence of wiring development processes from the connectome of adult *Drosophila*

**DOI:** 10.1101/2021.05.15.444029

**Authors:** Louis K. Scheffer

## Abstract

How is the brain wired during development? Here we look at some of the characteristics of the adult *Drosophila* brain, a result of this development process. From this we can speculate on at least some aspects of how the wiring was done. A common hypothesis is that surface proteins direct synapse formation among touching neurons. Assuming that surface proteins are specific to cell types, we find support for this hypothesis. The brain in general supports a wide variety of connections of different degrees of reciprocity, with ratios up to a thousand to one in the two directions of synapses between types. However, contacts between cells of the same type are always bi-directional and nearly equal strength. Furthermore, among similar cell types, at least in the mushroom body, the closer the cell type by morphology, the closer the inter-type contacts are to symmetrical. These findings are all consistent with the hypothesis that surface proteins specific to cell types determine the directivity of connections. Next we look at synapses per area, and find this varies widely and is roughly log-normally distributed. In most cases, the number of synapses saturates at higher areas, though other forms are seen - linear, flat, or decreasing with increasing area. Evidence suggests that at least some of this distinction is post-synaptic.

## 1 Introduction

The development of the adult brain, even in a relatively small animal such as *Drosophila*, is a complex and mysterious process. Mere tens of thousands of genes direct the development of circuits containing hundreds of thousands of neurons and hundreds of millions of synapses.

The connectome of the adult *Drosophila* shows only the results of this wiring process, which has completed long before the EM reconstruction. However, from the final circuits it may be possible to deduce some of the rules by which it was constructed.

One common theory of neural development holds that cells express interacting surface proteins that determine whether contact between two cells results in synapses[1][2], and if so, create a certain number of synapse per unit area[3].

If we make the additional assumption that cells of the same type express the same proteins, then whenever two cells of the same type touch, they should make the same number of synapses in each direction (refered to here as a symmetrical connection). This can potentially be examined in a dense connectome, in which every cell is typed and all synapses are annotated. Such a connectome can also be used to examine how synapse counts vary with contact are among the connections between different cell types.

## 2 Methods

We use the recent dense connectome containing roughly half of the brain (a hemibrain) of an adult female *Drosophila*[4]. This connectome, including cell names, cell types, connections, and synapse counts is available at https://neuprint.janelia.org. Version 1.2 of the connectome was used in this analysis. We query this connectome using code that can be found at https://github.com/janelia-flyem/SameTypeAnalysis. For the second portion of our analysis, an additional file is needed containing the cell to cell contact areas. This file (for the 1.2 version of the connectome) is also included in the GitHub repository.

In general, we simplify our analysis by looking only at each connections in terms of the types of the partners. The hemibrain connectome contains about 25,000 cells, but only about 5000 cell types. In the first part of this study we look at type to type connectivity. This matrix is sparse, with about 1.1 million non-zero entries of about 25 million total, as each type typically forms synapses with about 220 other types.

For our analysis, we first look at the connections between all types, showing that biology has no difficulty constructing very asymmetrical connections. We then look at connections between cells of the same type, and check if they are more symmetrical, as hypothesised. Finally we look at a set of morphologically similar cells, the Kenyon cells, looking to see if cells that are closer matches morphologically are also closer to symmetrical.

In a second part of this analysis, we examine how the number of synapses depends on the area of contact. Given the area and the synapse count, we then find Pearson’s correlation coefficient between the two, fit functions to the resulting data, and classify the connections into different types.

## 3 Results

### 3.1 Connections between cells of the same type

Connections in the *Drosophila* brain show a wide variety of directionality. For any pair of cell types, the contact area between the types is the same in either direction (*A* → *B* or *B* → *A*) but the number of synapses is not. Fig. 1 shows the range of connectivities.

**Figure 1:**
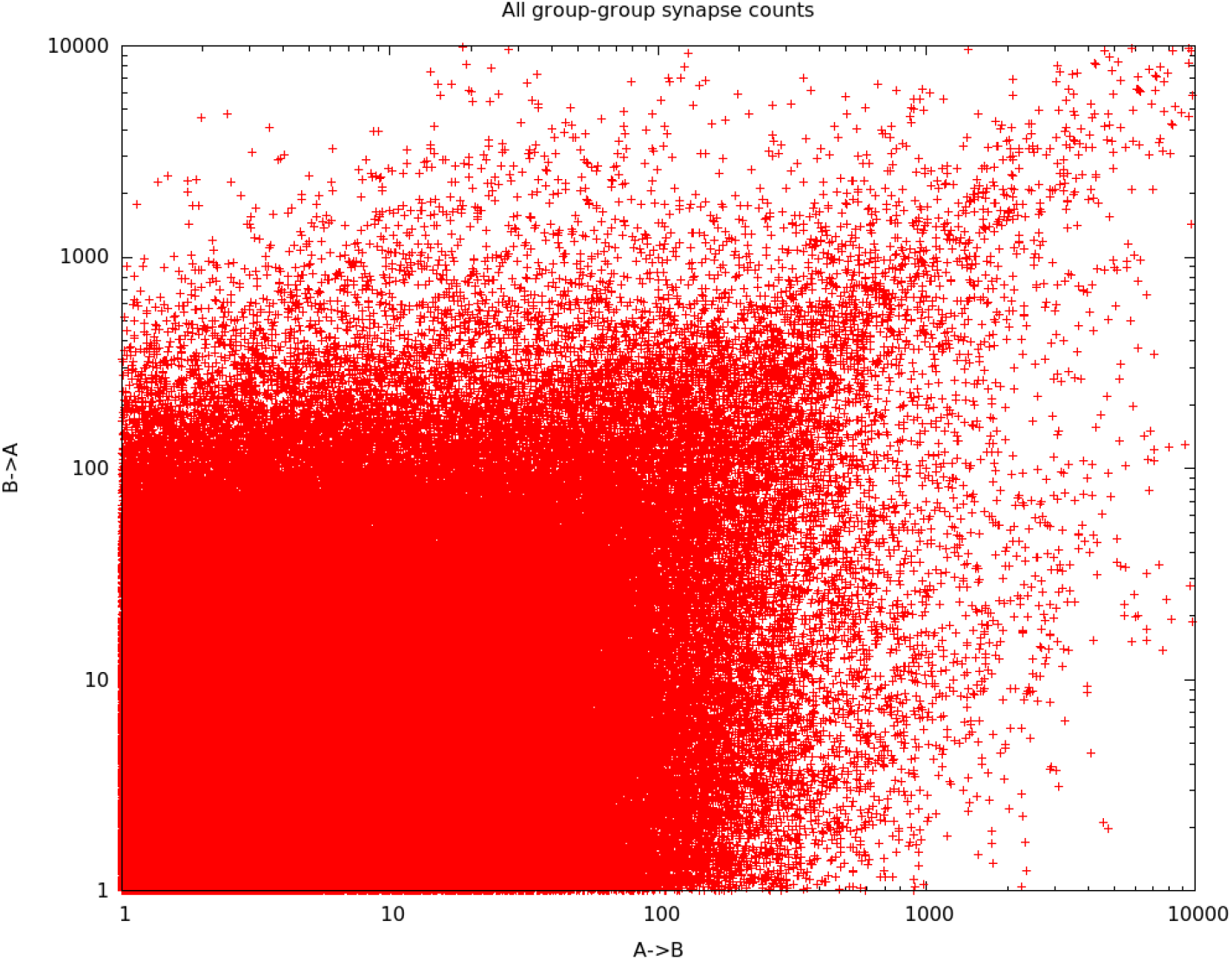
Type to type synapse counts, showing that biology supports a wide range of reciprocity. Roughly 1.7 million connections are shown. Points are “fuzzed” by +− 0.5 to show density (rather than points) at small integer values.

There are a significant number of connections with very high directionality. Table 1 shows connections with more than a thousand synapses in one direction and ln(ratio of synapses) > 7, or roughly > 1100: 1. These are divided into three groups - the optic lobes, the central complex, and the mushroom body.

**Table 1:**
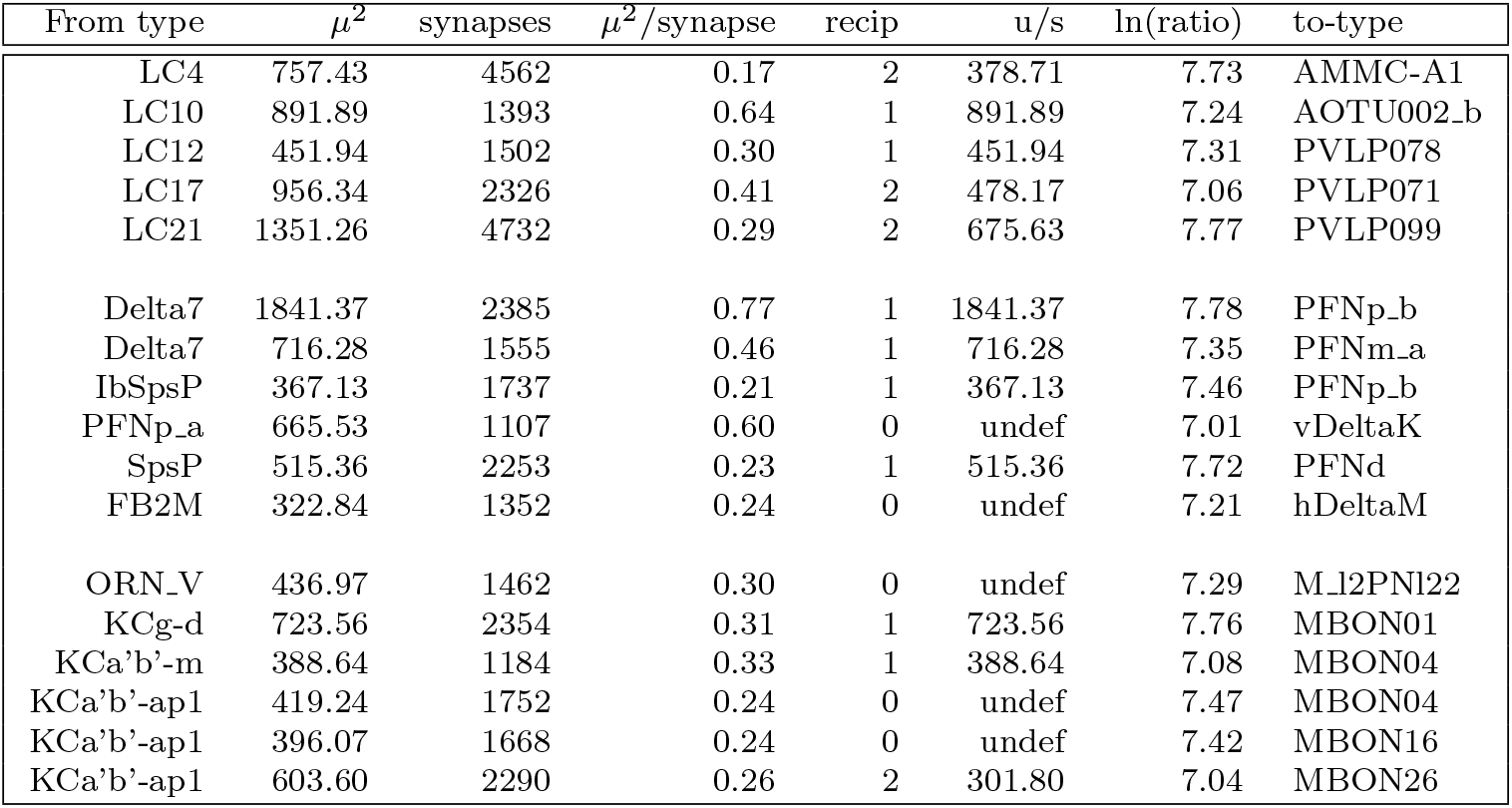
Type to type connections with extreme directionality of synapses, with a ratio of more than 1000:1. Three groups are shown - the optic lobes, the central complex, and the mushroom body. Zero values are replaced by 1 when calculating ratios, which would otherwise be infinite.

Why are all the examples in the well studied regions? There are several possibilities. First, the requirement for 10,000 synapses limits the examples to regions with many repeating types. Other parts of the brain have less repetition. There may also be a selection effect. Synapse detection gives a small fraction of false positives. In these well studied regions most of these have been eliminated by hand editing. In other regions more may exist and prevent the directionality from reaching the 1000:1 threshold.

So biology is perfectly capable of creating connections that are very asymmetrical, with many more connections *A* → *B* than *B* → *A*. However, if this asymmetry is created by a mechanism that depends on the proteins expressed by the touching cells, then there should be cases where this asymmetry is not possible. This would be when a cell touches a cell of the same type, assuming that all cells of the same type express similar proteins. The expressed surface protein theory would then predict that all such connections are symmetrical, with roughly equal number of synapses in both directions.

We can test this prediction using the dense connectome of the hemibrain[4]. This connectome contains 22 different cell types that have at least 10,000 synapses to other cells of the same type. These types are shown in Table 2.

**Table 2:**
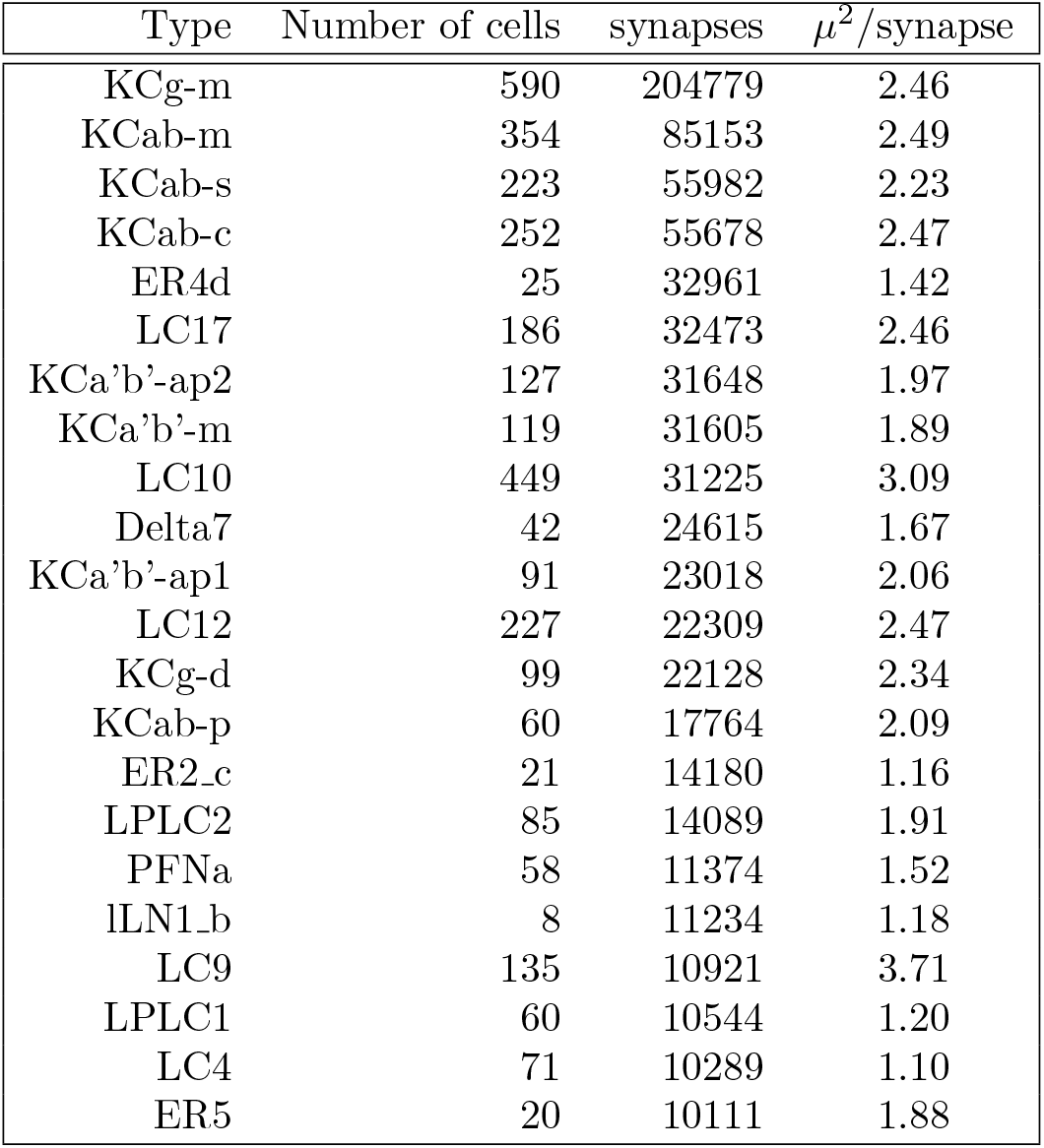
Cells in hemibrain data set with more than 10,000 synapses between cells of the same type.

In this case we cannot simply look at the type to type connectivity, since by definition *A* → *B* and *B* → *A* have exactly the same synapse count if *A* = *B*. Therefore if one attempts to plot same-type to same-type connections in the same way as Fig. 1, it will yield only a single row of points along the diagonal. Instead we need to look more closely, at the individual connections between neurons of the same type. These are shown in Fig. 2 on a linear scale, and in Fig. 3 on a log scale.

**Figure 2:**
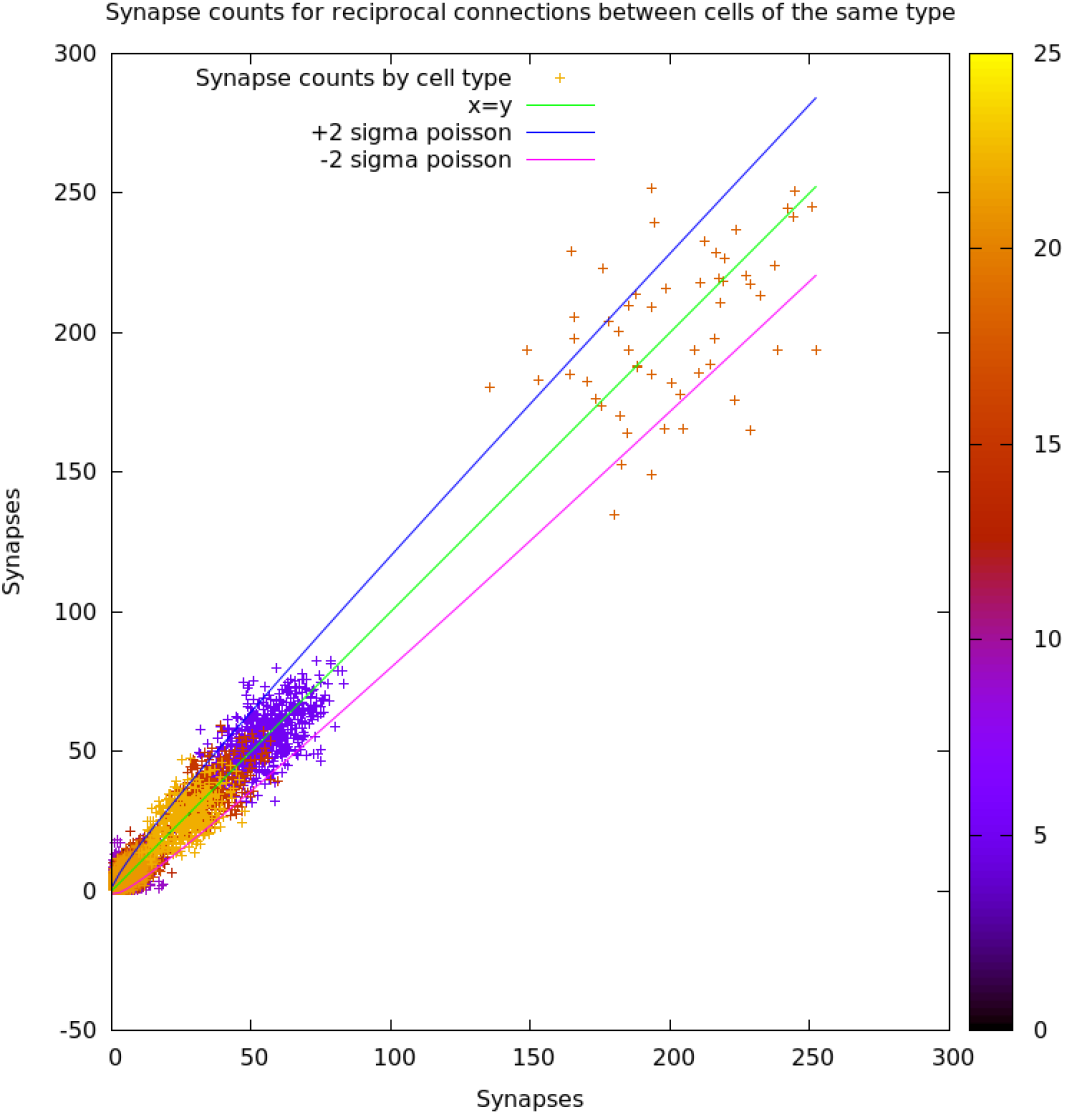
Synapse counts between reciprocal connections between cells of the same type. Color of point corresponds to the scale on the right, which in turn corresponds to position in the table of cell types above. The plot is of necessity symmetrical about y=x since each connection can be viewed from either side. Lines labelled ‘poisson’ show the range that would be expected if the creation process was strictly Poisson.

**Figure 3:**
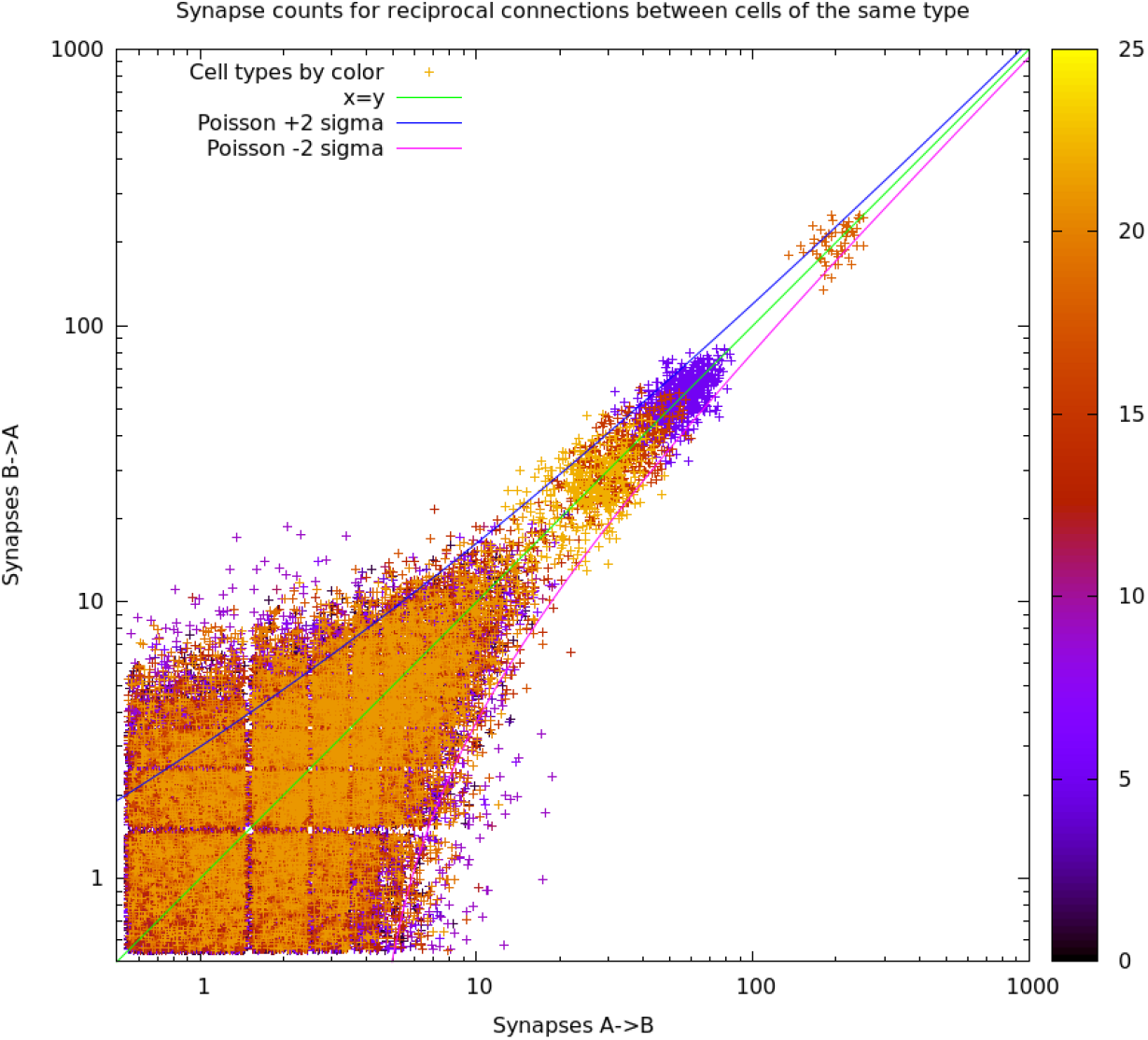
Plot for connections between cells of the same type. Roughly 98000 connections from 22 cell types are shown. Point color corresponds to the scale on the right, which in turn corresponds to the table of cells above. Points are fuzzed to show density.

### 3.2 Nearly same type connections

The above analysis looks at connections between cells of the same type. We can expand this by looking at connections between cells of different type. If the surface protein hypothesis is correct, we would expect that the more closely related the cell type, the more closely the connections would be to symmetrical. This can be tested with Kenyon cells[6], which are composed of closely related but not identical types. In the hemibrain data, we identify 14 Kenyon cell types. Four of them, the KC-gamma-s types, consist of only one cell so we lump the 4 cells together here, leaving 11 sub-types.

Which types are “most similar” (at least based on morphology) is somewhat subjective. A likely grouping would be the alpha-beta-prime KCs (KCa’b’-ap1, KCa’b’-ap2, and KCa’b’-m) as one group, the main alpha lobe cells as another (KCab-c, KCab-m, and KCab-s), the alpha posterior cells (KCab-p) as a third, and all the gamma lobe KC cells (KCg-d, KCg-m, KCg-t, and KCg-s.*) as the fourth.

We count the number of synapses between these types, as shown in Table 3.

**Table 3:** KC to KC type synapse counts. The first entry in each row is the ‘from’ cell type, and the column indicates the ‘to’ cell type. For brevity, all type names omit the starting “KC”, so (for example) the column “ab-s” refers to type “KCab-s” in the hemibrain data.

|  | a'b'-ap1 | a'b'-ap2 | a'b'-m | ab-c | ab-m | ab-s | ab-p | g-d | g-m | g-t | g-s.* |
| --- | --- | --- | --- | --- | --- | --- | --- | --- | --- | --- | --- |
| a'b'-ap1 | 23018 | 2089 | 54 | 22 | 22 | 35 | 4 | 29 | 953 | 9 | 24 |
| a'b'-ap2 | 1932 | 31648 | 6208 | 53 | 119 | 86 | 22 | 13 | 649 | 6 | 14 |
| a'b'-m | 57 | 6110 | 31605 | 111 | 205 | 155 | 226 | 2 | 285 | 0 | 1 |
| ab-c | 68 | 317 | 484 | 55678 | 12310 | 665 | 27 | 14 | 1202 | 5 | 2 |
| ab-m | 41 | 243 | 514 | 12009 | 85153 | 12572 | 120 | 12 | 873 | 4 | 1 |
| ab-s | 68 | 237 | 432 | 604 | 13168 | 55982 | 3356 | 10 | 1298 | 6 | 2 |
| ab-p | 15 | 50 | 248 | 10 | 94 | 2474 | 17764 | 3 | 82 | 0 | 31 |
| g-d | 15 | 11 | 1 | 2 | 0 | 2 | 2 | 22128 | 7098 | 697 | 1279 |
| g-m | 963 | 641 | 291 | 384 | 509 | 622 | 50 | 6141 | 204779 | 1627 | 1218 |
| g-t | 10 | 3 | 0 | 3 | 2 | 3 | 0 | 691 | 1687 | 152 | 51 |
| g-s.* | 30 | 7 | 3 | 0 | 1 | 2 | 20 | 1226 | 1391 | 57 | 145 |

We can also express these synapse counts as the ratio of the synapse count to that in the opposite direction. Almost all of these have ratios of less than 4:1, much less than biology is capable of (see Fig. 1). This is shown in Table 4.

**Table 4:**
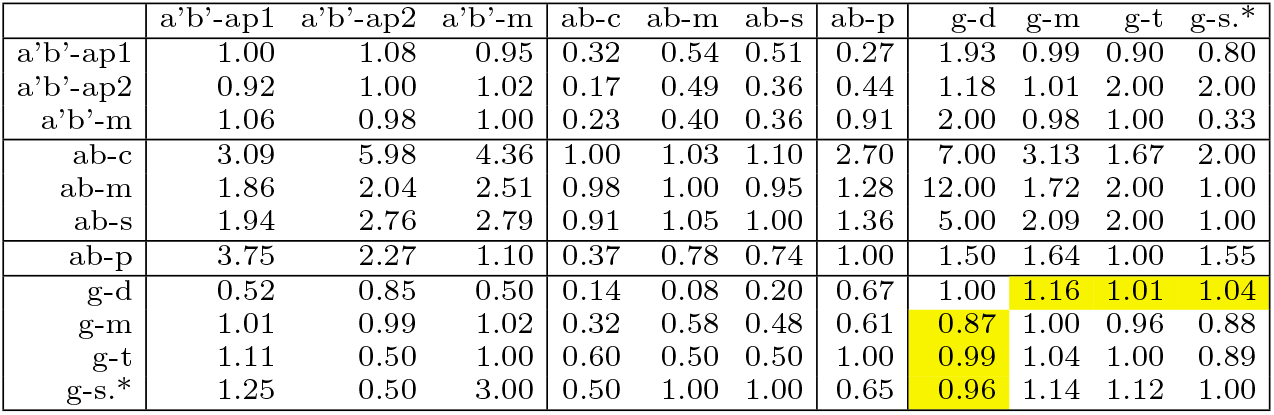
KC to KC type connections expressed as ratios of the counts in both directions. For brevity, all type names omit the starting “KC”, so (for example) the column “ab-s” refers to type “KCab-s” in the hemibrain data. The shaded squares indicate interactions where one type receives olfactory input and the other visual input.

To see if the more closely related groups have more symmetrical connections, we can average these ratios over the purported groups. This is shown in Table 5. the results are consistent with the surface protein hypothesis. None of the ratios are large, reflecting the assumption that these cells are all closely related. Within each group, where the cells are most similar, the connections are very close to symmetrical. Between groups, the largest differences are found between KCgamma and KCab, and between KCa’b’ and KCab. The most similar groups are KCa’b’ and KCgamma.

**Table 5:**
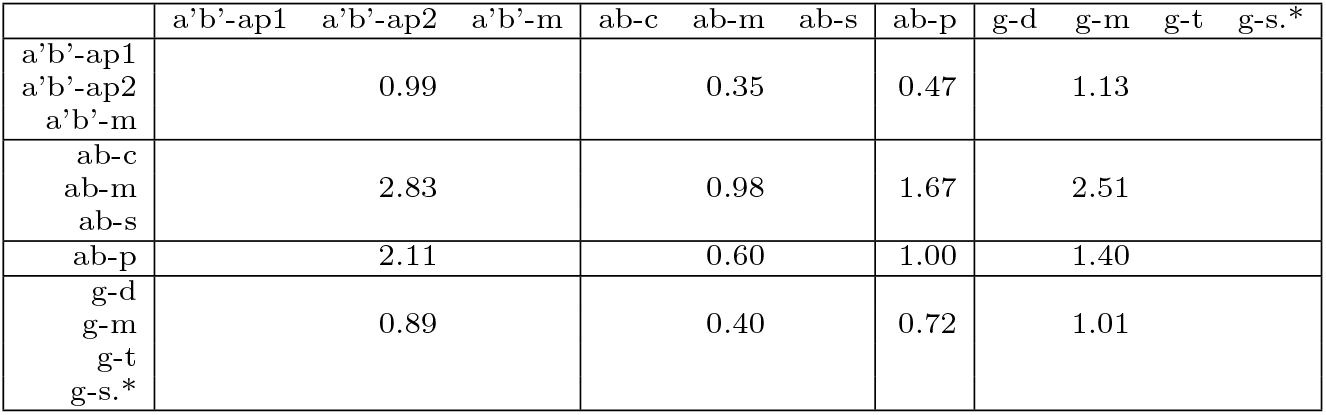
Summary of geometric mean of the ratios of the off-diagonal elements. In the diagonal boxes, only elements above the diagonal are used. The entry for KCab-p →KCab-p is defined as 1.00 since there are no off-diagonal elements to average. For brevity, all type names omit the starting “KC”, so (for example) the column “ab-s” refers to type “KCab-s” in the hemibrain data.

This could lead to a suggested dendrogram as shown in Fig. 4. It would be very interesting to compare this analysis to a dendrogram computed from gene expression analysis. The relatively smooth dropoff as types become more different implies a mechanism that can generate such “analog” reponses, as opposed to a strictly ‘connect or not’ response.

**Figure 4:**
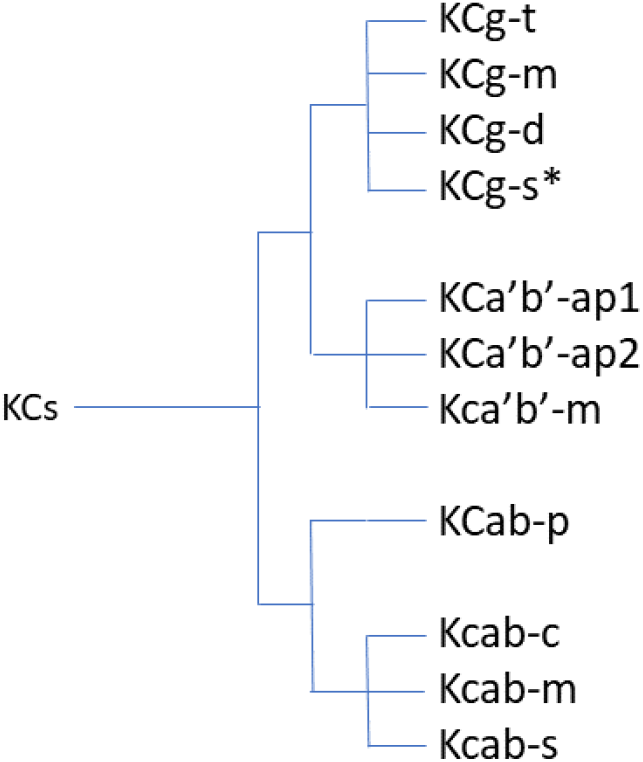
Dendrogram based on reciprocity among synapse counts. The position of KCab-p is not completely certain. The other groupings are strongly suggested by the data.

An alternative (or an addition) to the surface protein specification is synapse contruction based on activity[5]. Between cells of similar type, this could be hard to distiguish from the surface protein model, since cells of a similar type might well have similar activity patterns. However, the *Drosophila* mushroom body offers a potential test. The KCg-d neurons get primarily optic input, where the other KG-g neurons recieve primarily olfactory input[6]. Presumably the input patterns might be different in these cases, perhaps because visual input changes more quickly than the olfactory environment. However, the connections between the KGg-d and other KCg neurons are consistent with all the other KC-g connections, where both members of each pair recieve olfactory input, as shown in Table 4. This provides circumstantial evidence against the activity based model.

### 3.3 Synapse count as a function of area

The surface protein hypothesis says that all connections between cells of the same type are symmetrical, and seems supported by the data above. However, symmetry does not depend on how synapse count varies as a function of area, since the connection can still be symmetrical even if the synapse numbers in each direction are a strong function of area. So analyzing synapse counts per unit area provides another window into possible synapse development strategies.

To analyze this further, we calculated the synapse count versus area for all connections in the hemibrain. In general, increased area leads to increased synapse count. We measured this by computing the correlation between area and synapse count, which is generally positive. Over the whole brain, the correlation between area and synapse count is typically 0.6 to 0.9, as measured by Pearson’s r. This is shown in Fig. 5.

**Figure 5:**
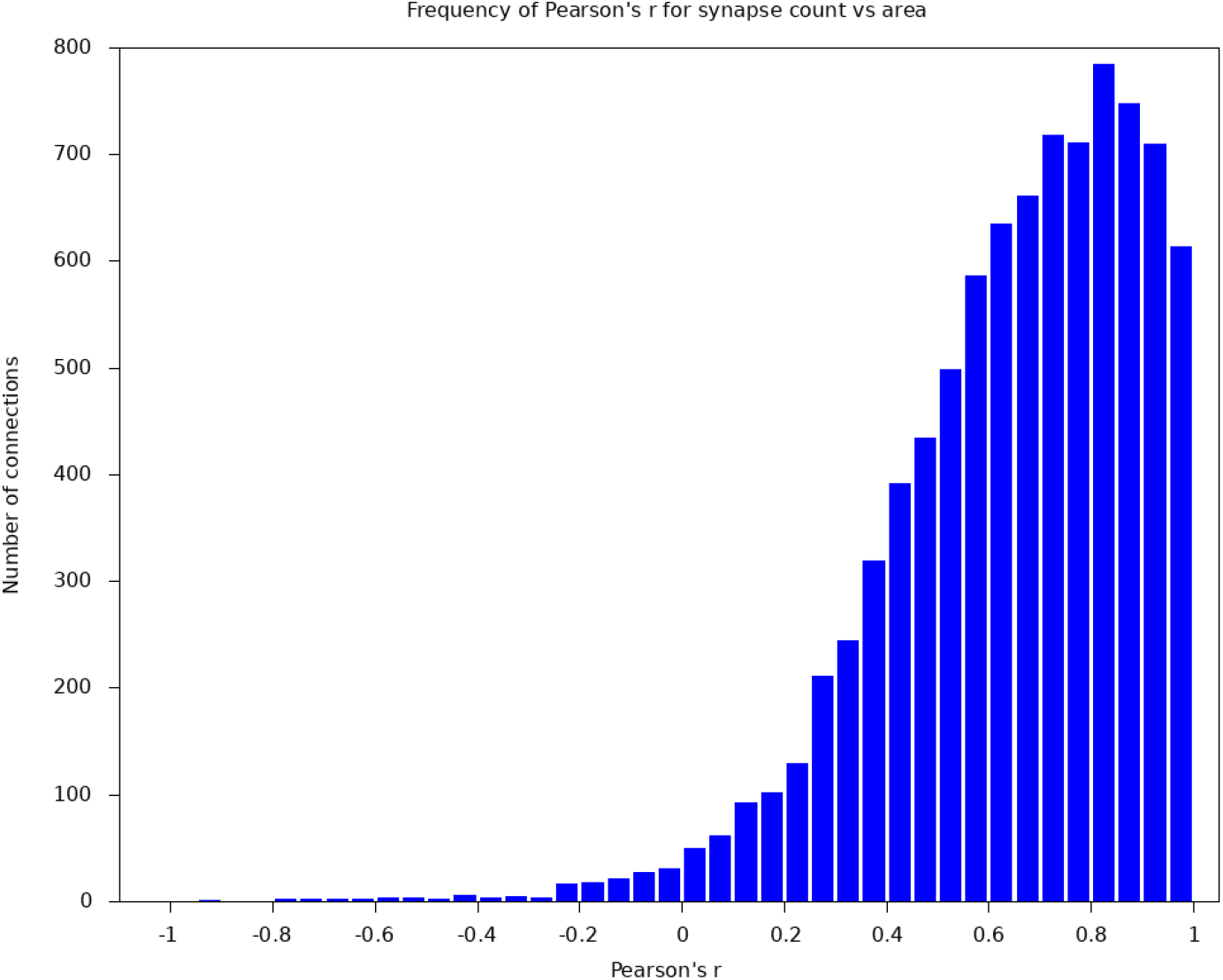
Histogram of Pearson’s r values for the 1150 type-to-type connections with at least 1000 synapses.

However, even with the same correlation, different cell types could have a different number of synapses per unit area. For example, this seems true of the connections between the two cell types ER2_c and ER5, as shown in Fig. 6.

**Figure 6:**
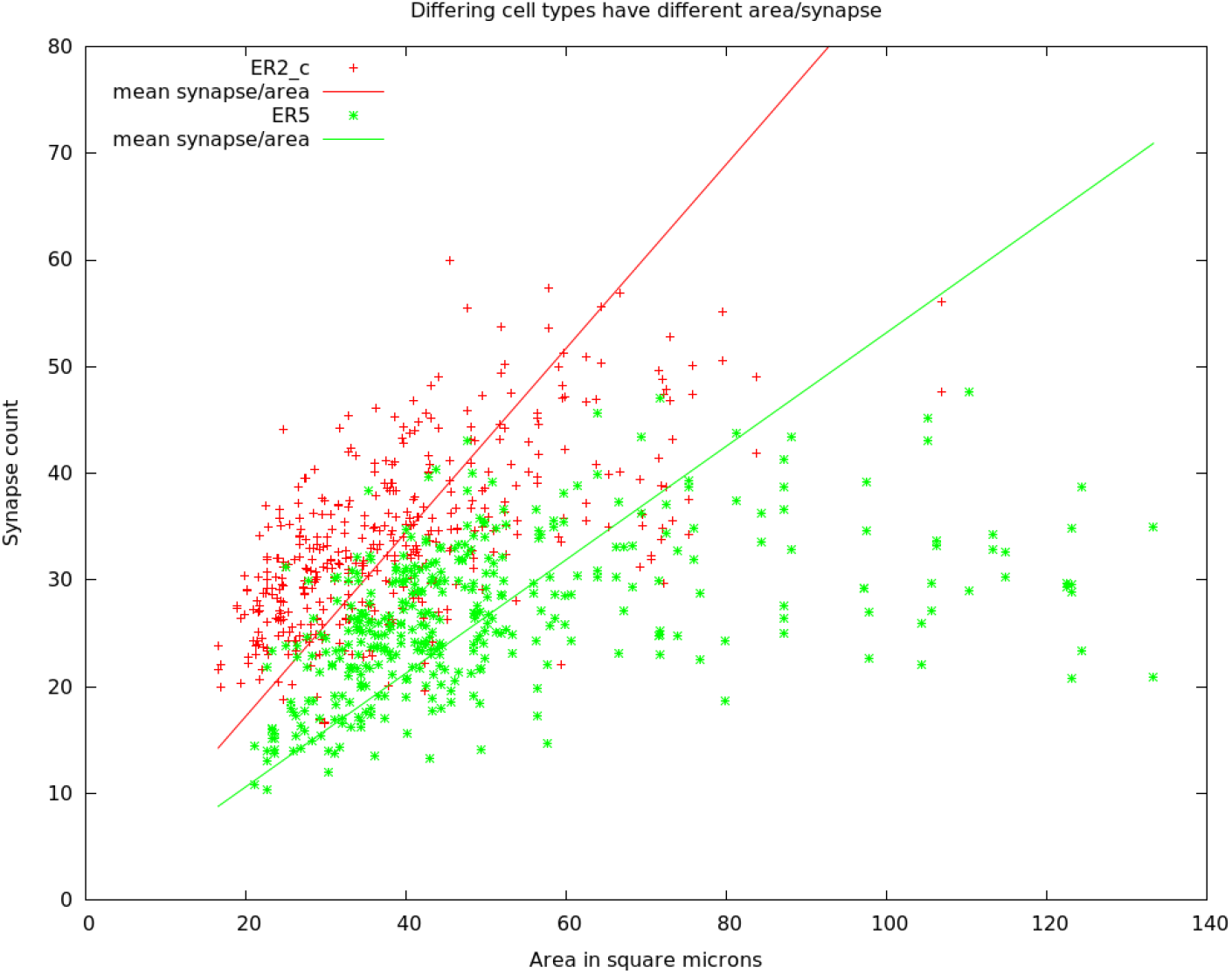
Area and synapse count for within-type connections for two cell types. Most cells have a synapse count that is roughly proportional to area, with a tail having higher areas but similar synapse count. The solid lines show the predicted number of synapses for each type, given the mean square microns/synapse of each type.

In general, the synapse count per area covers a wide range, from about 0.05-20 square microns per synapse. The area/synapse is rougly log-normally distributed, as shown in Fig. 7. The peak is at roughly 0.5 square microns per synapse.

**Figure 7:**
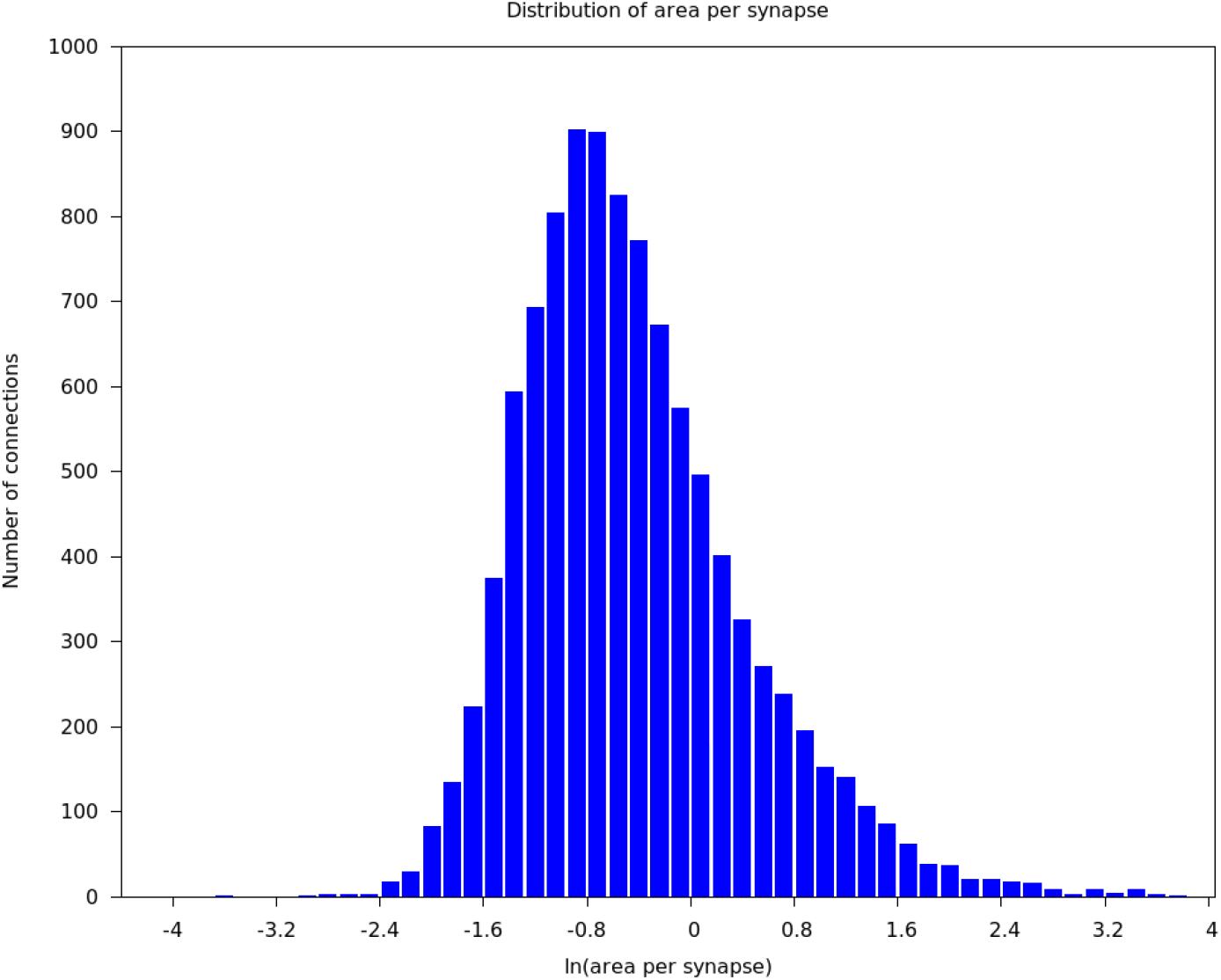
Histogram of the area/synapse over all type-type connections with at least 100 synapses.

Next, we observe the connections between cells of the same or different types is not always strictly proportional to the area of contact. We qualitatively divide these into several behaviors. The most common, after proportional, is what we call “Saturated”, where the number of synapses no longer goes up once the area reaches a certain limit. We model this (strictly empirically) with a logistic function. In addition, less commonly we see cases where the correlation between area and synapse count is near zero, or even negative.

An example where the correlation is very high, and the synapse count is well described by a linear function, is shown in Fig. 8.

**Figure 8:**
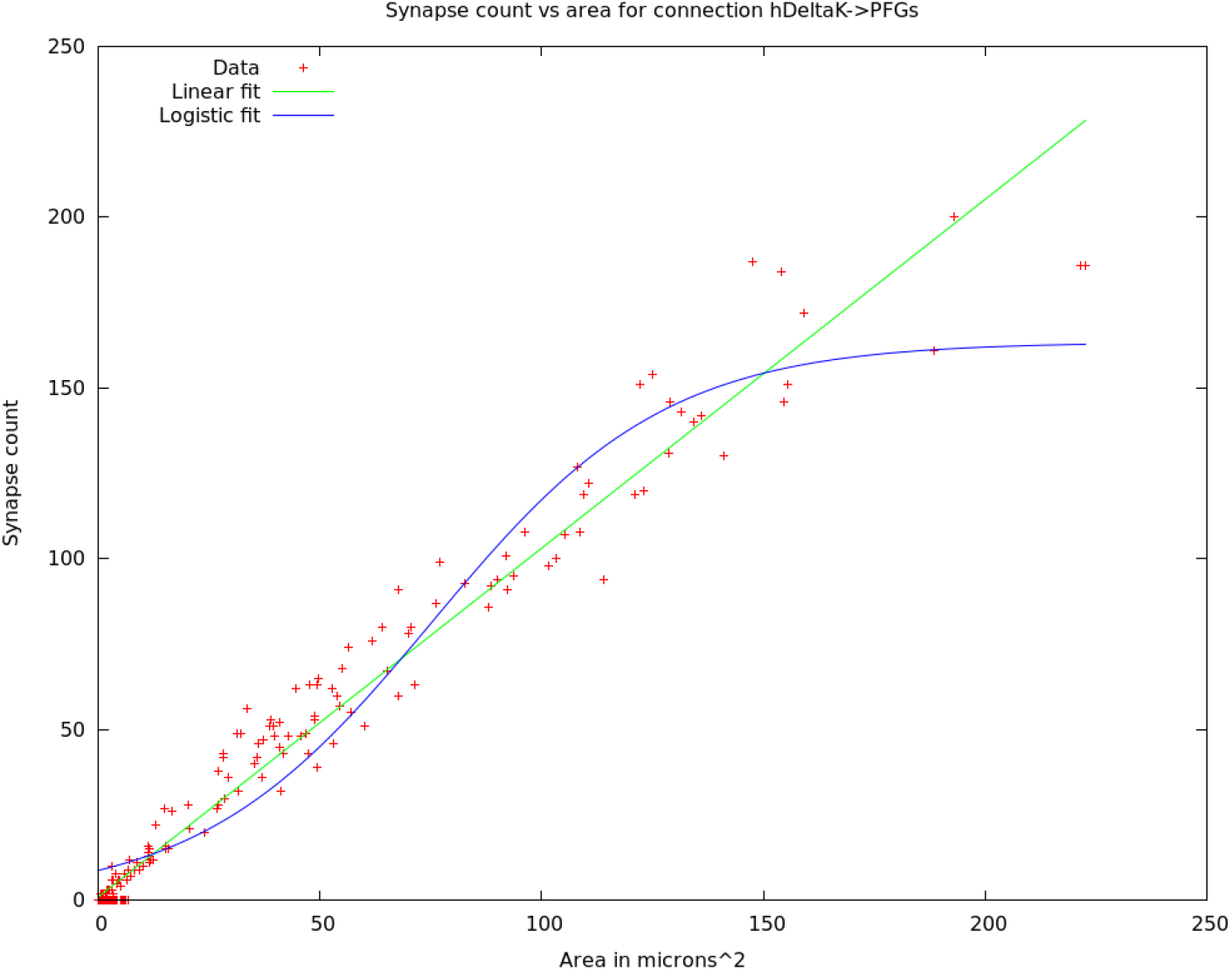
A case where the synapse count is well correlated with area. In this examples, Pearson’s r > 0.9, and a linear fit has much smaller residuals than a logistic fit.

However, not all connections are well fit by a linear function, as shown by the tail of the synapse count of ER5 in Fig. 6, where the synapse count levels off after a certain area is reached. These connections are better fit with a logistic function, as shown in Fig. 9.

**Figure 9:**
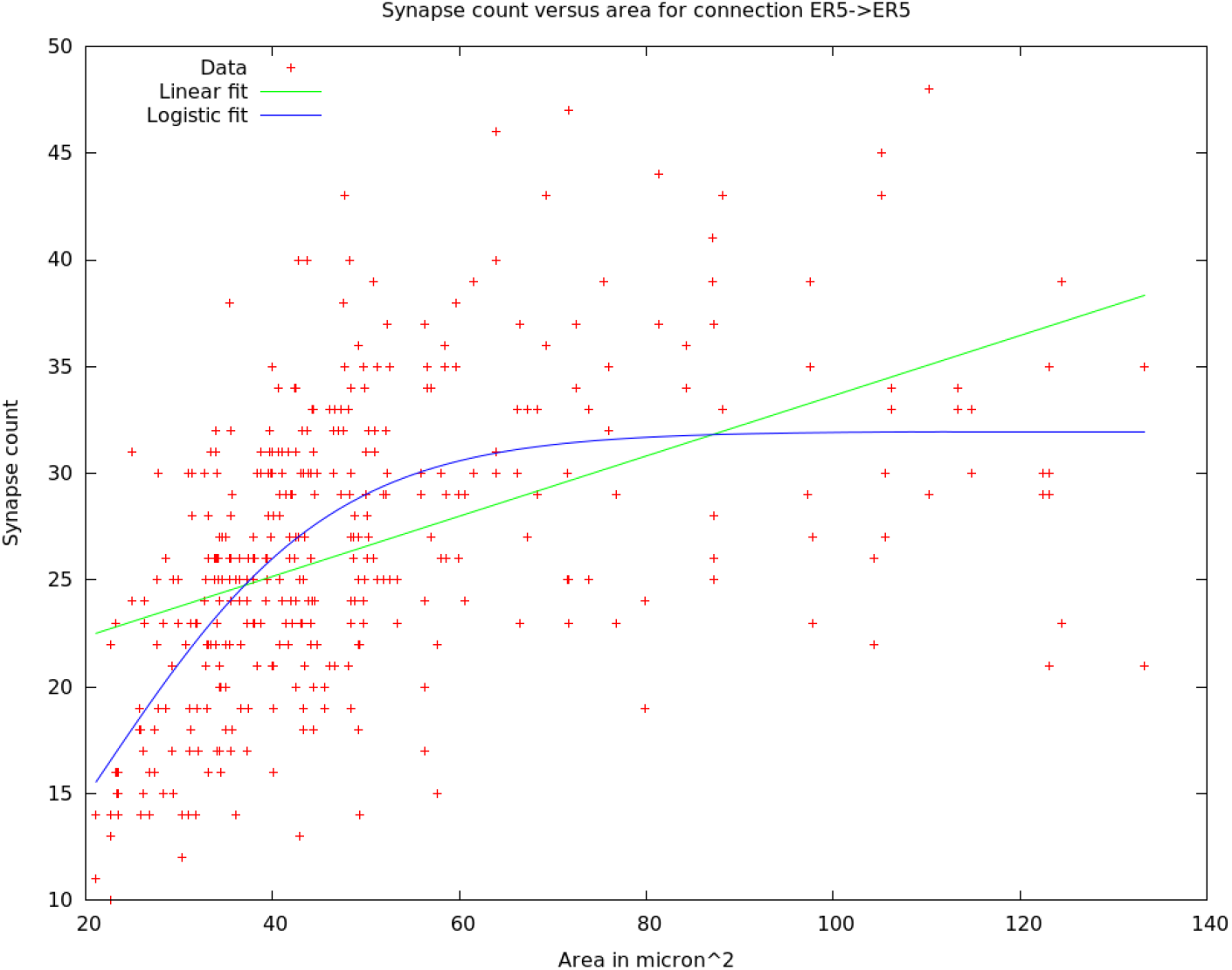
A case where the synapse count is more closely described by logistic function, as opposed to a linear function.

Next, there are some connections that have no overall correlation with area, as shown in Fig. 10. Some other mechanism must determine the synapse count in these cases.

**Figure 10:**
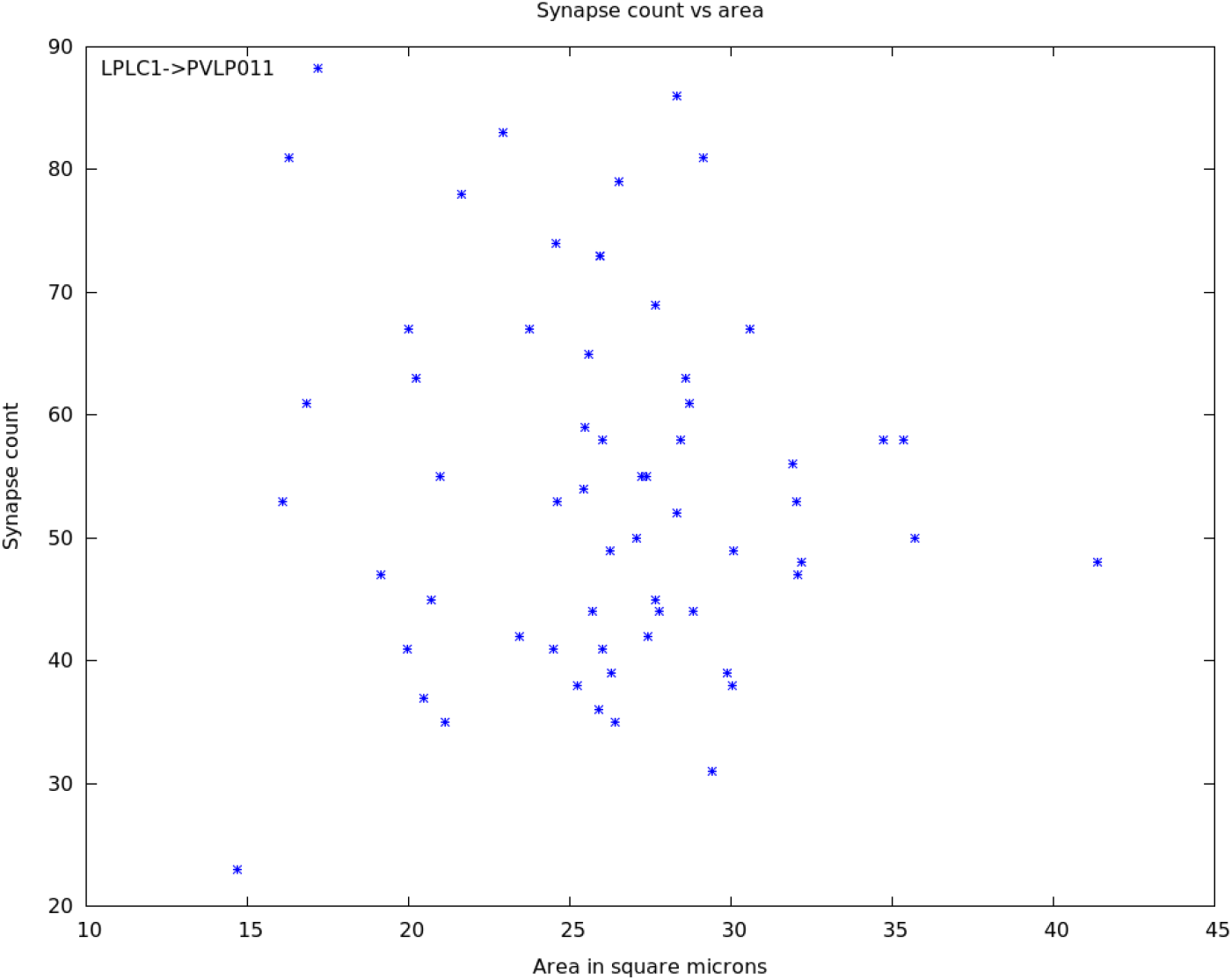
A case where there is essentially no correlation between contact area and synapse count. Pearson’s *r* ≈ − 0.02

Finally, there are rare cases where the correlation is negative - cells with larger contact area form fewer synapses on the average. The mechanism behind this remains mysterious. Two of these connections are shown in Fig. 11. This connection was chosen for plotting since the three connections shown share the same source type (lLN2T_a), and very similar target types (lLN1_a, lLN1_b, and lLN1_c). Here the ba’, ‘_b’, and ‘_c’ indicate connectivity subtypes, or cells that have very similar morphology but differ in their connectivity[4]. As shown in the graph, the ba’ subtype has the usual positive slope, whereas ‘_b’ and ‘_c’ are negative. This implies the area/synapse relationship is determined, at least in part, on the post-synaptic side.

**Figure 11:**
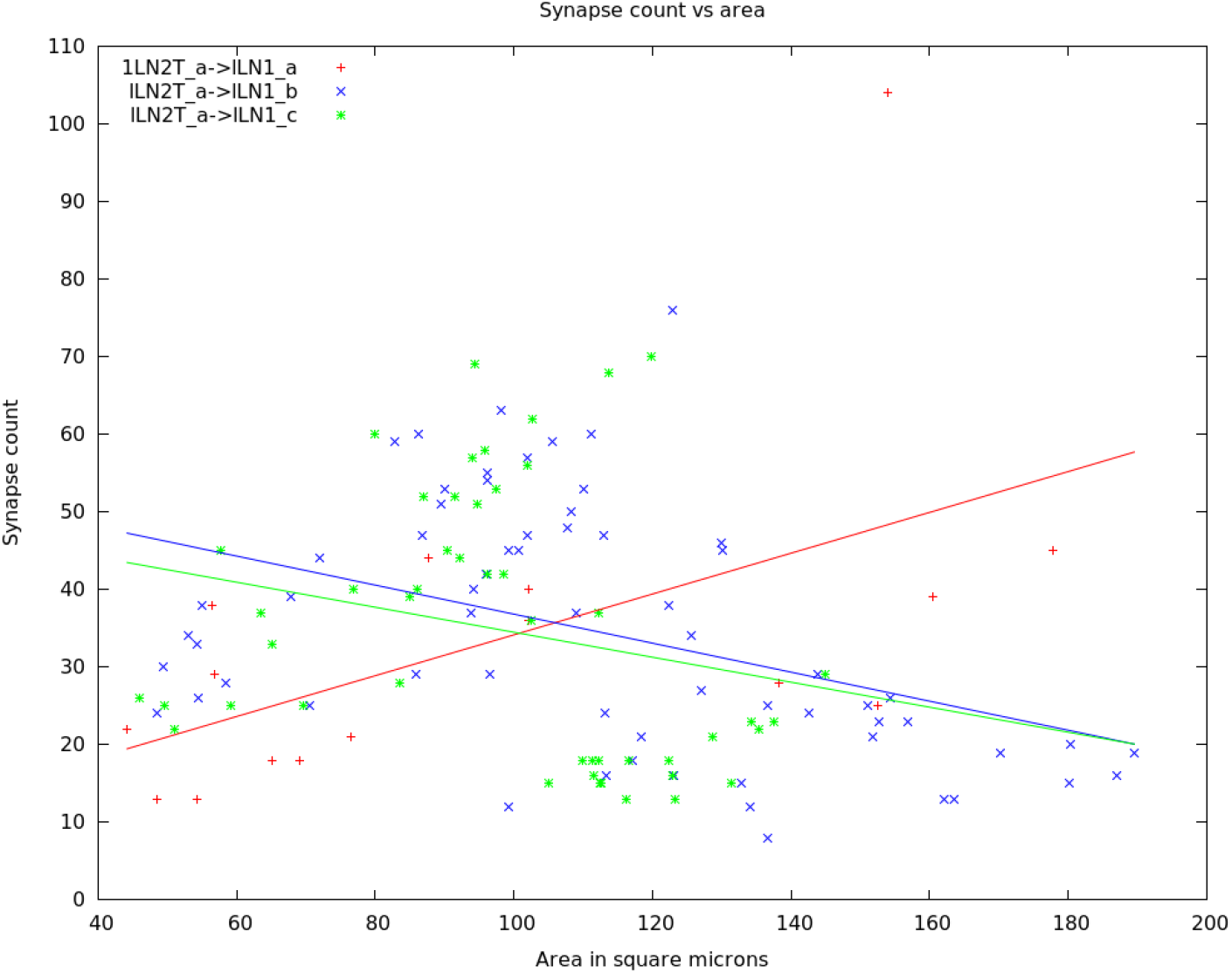
Two cases where there is a negative correlation between contact area and synapse count, and one case with a positive correlation. Pearson’s *r* ≈ −0.2 for the decreasing cases, and a linear fit shows negative slopes. All three cases share the same source type, and differ only in the connectivity subtypes of the target type.

Which functional form of area/synapse count is most common? For each connection with enough data (here at least 100 synapses and at least 5 non-zero synapse counts), we first look at correlation, and then fit both a linear and a logistic function to the synapse vs. area curve. We then classify these somewhat arbitrarily. If the correlation is < −0.1, we say the curve is ‘Falling’, which accounts for 0.8% of the connections. Next is if the correlation is −0.1 < *r* < 0.1, which we call ‘Flat’, and accounts for 1.2%. Otherwise the function is rising, and we call it ‘Linear’ (46%) or ‘Saturated’ (51%) depending on which gives the better fit (compensating for the 3 parameters of the logistic model versus 2 parameters of the linear model).

## 4 Conclusions

The cell surface model is supported by the observation that connections between same-type cells are largely symmetrical, and that among similar-type cells, the more similar the morphological type, the more nearly the connection is symmetrical. This holds even when the two types receive different sensory modalities.

The synapse/area between cells varies widely, and is roughly log-normally distributed. In addition, another mechanism must also be in play to account for the occasional neutral or negative dependence of synapses on area. Examples show this is determined, at least partially, on the post-synaptic side.

## Notes

https://neuprint.janelia.org

https://github.com/janelia-flyem/SameTypeAnalysis

### Competing Interest Statement

The authors have declared no competing interest.

## References

[1] Matthijs Verhage, Ascanio S Maia, Jaap J Plomp, Arjen B Brussaard, Joost H Heeroma, Hendrika Vermeer, Ruud F Toonen, Robert E Hammer, Markus Missler, Hans J Geuze, et al. Synaptic assembly of the brain in the absence of neurotransmitter secretion. Science, 287(5454):864–869, 2000.

[2] Kang Shen. Molecular mechanisms of target specificity during synapse formation. Current opinion in neurobiology, 14(1):83–88, 2004.

[3] Kang Shen and Peter Scheiffele. Genetics and cell biology of building specific synaptic connectivity. Annual review of neuroscience, 33:473–507, 2010.

[4] Louis K Scheffer, C Shan Xu, Michal Januszewski, Zhiyuan Lu, Shin-ya Takemura, Kenneth J Hayworth, Gary B Huang, Kazunori Shinomiya, Jeremy Maitlin-Shepard, Stuart Berg, et al. A connectome and analysis of the adult *Drosophila* central brain. Elife, 9:e57443, 2020.

[5] Larry C Katz and Carla J Shatz. Synaptic activity and the construction of cortical circuits. Science, 274(5290):1133–1138, 1996.

[6] Feng Li, Jack W Lindsey, Elizabeth C Marin, Nils Otto, Marisa Dreher, Georgia Dempsey, Ildiko Stark, Alexander S Bates, Markus William Pleijzier, Philipp Schlegel, et al. The connectome of the adult *Drosophila* mushroom body provides insights into function. Elife, 9:e62576, 2020.

